# Investigating Increased CO_2_ concentration on the pH of various plant species

**DOI:** 10.1101/2021.11.13.468474

**Authors:** Joshua Schafer, Troy Puga, Pearce Harris, Nora Strasser, Gary Branum, Prince N Agbedanu

**Affiliations:** Department of Health Sciences, Friends University, Wichita Kansas 67213, USA

## Abstract

The concept of bioremediation is quickly becoming the norm in the resolution of environmental issues. The steady increase in carbon dioxide (CO_2_) levels, as documented by NASA, inspired scientists to engineer plants to absorb excess CO_2_ from the atmosphere. Here, we have explored the consequences of the uptake of excess CO_2_ by select plants. Carbon dioxide dissolves in H_2_O to produce H_2_CO_3,_ which dissociates to yield H^+^ ions. We hypothesized that increased CO_2_ absorption results in decrease in pH of plant sap. Three plants (*Byophyllum pinnatum*, Romaine Lettuce and Nevada Lettuce), exposed to increased CO_2_ concentrations (15%), demonstrated a consistent increase in pH towards alkalinity compared to control plants. Based on the outcome being opposite of what we have hypothesized, our results suggest *Byophyllum pinnatum,* Romaine lettuce and Nevada lettuce, all have a unique homeostatic system to prevent over-absorption of CO_2_ in a CO_2_-rich environment.

## Introduction

This work is inspired by the concept of bioremediation; the use of naturally occurring or genetically engineered organisms to solve environmental problems. Carbon dioxide levels have steadily increased over the years, raising concerns for individuals who want to solve this problem. In Scientists quests to address this problem, the idea of engineering plants to increase the capacity of CO_2_ absorption from the environment is still in the early stages. Others have pursued the use of a semi-synthetic rubisco, which has the ability to increase the rate of photosynthesis [2]. We ask the question, what will be the effect of excess CO_2_ on the pH of plant sap? Theoretically, carbon dioxide dissolves in water to produce carbonic acid according to equation [1].

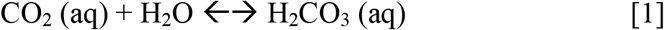

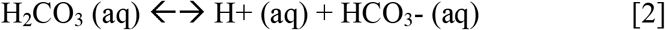

According to equation [2], dissolved CO_2_ in the form of H_2_CO_3_ may lose up to two protons through the acid equilibrium. The relatively small amounts of H+ produced, when built up, is anticipated to decrease the pH of plant cytosol. We hypothesize that the exposure of plants to increased CO_2_ levels results in a decrease in the pH of plant cytosol.

### Objectives

- Monitor pH of plant sap at increased CO_2_ concentrations
- Determine whether bioremediation is a safe solution to combat CO_2_ pollution based on pH data
- Further understand the relationship between CO_2_ and pH of plant sap

### Materials

- pH meter
- Garlic press
- CO_2_ incubator
- Lamp for light source
- Plant soil heater

### Method

Plants (e.g. *Byophyllum pinnatum*) were incubated in the presence of increased CO_2_ with a light source or under normal atmospheric conditions with a light source. The lighting schedule, temperature and watering of the two plants were kept the same, with the only variable being the difference in CO_2_ concentration. The control plant was maintained in atmospheric CO_2_ concentrations and the test plant was maintained in 15% CO_2_ concentration. The light sources were on for approximately ten hours per day and the plants were watered once per week. The same set up was observed for 6 stalks each of *Romaine lettuce* and *Nevada lettuce*. Every 2 to 3 days, leaves from each plant was homogenized for sap extraction and pH testing. The results are as follows:

### Results: *Byophyllum pinnatum*

**Table 1:** pH readings vs exposure time in *Byophyllum pinnatum*

| Day | Control (pH) | Test (pH) |
| --- | --- | --- |
| 3 | 4.34 | 4.98 |
| 5 | 4.32 | 4.87 |
| 7 | 4.54 | 5.32 |
| 11 | 3.97 | 4.57 |
| 12 | 4.88 | 5.22 |

### Results: *Byophyllum Pinnatum*

**Figure 1:**
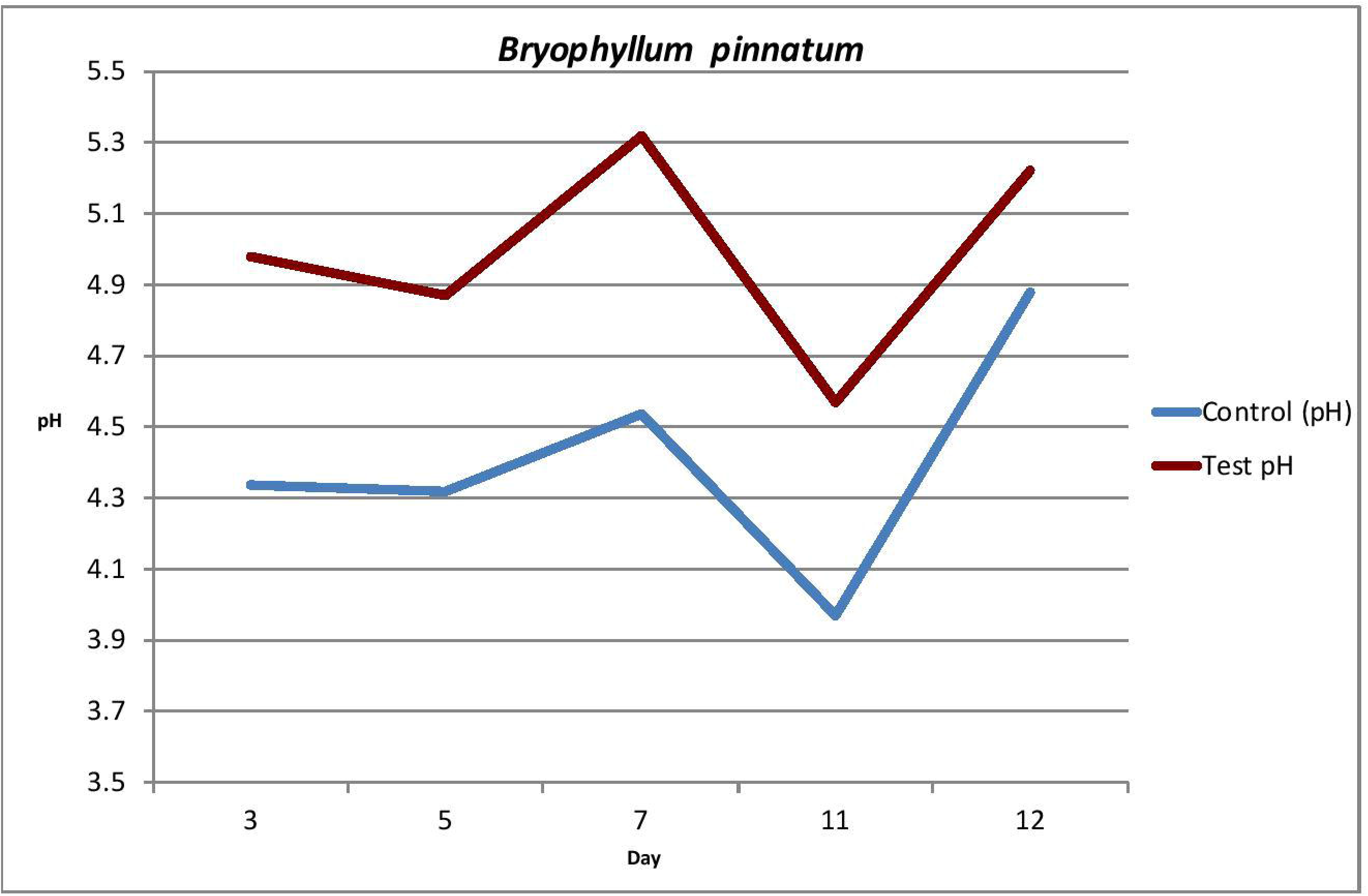
The results were not what we expected. The plant exposed to excess CO_2_ became more alkaline instead of more acidic, while the control plant remained well under pH=5. This suggests *Bryophyllum pinnatum* has a homeostatic system to prevent over-absorption of CO_2_ in a CO_2_-rich environment.

### Results: *Romaine lettuce (Lactuca sativa, variety longifolia)*

**Table 2:** pH readings vs exposure time (Days) in Romaine lettuce

| Day | Control<br>(pH) | Test (pH) |
| --- | --- | --- |
| 4 | 5.78 | 5.9 |
| 7 | 5.63 | 6.48 |
| 11 | 6.21 | 6.58 |
| 14 | 6.03 | 6.75 |

**Figure 2:**
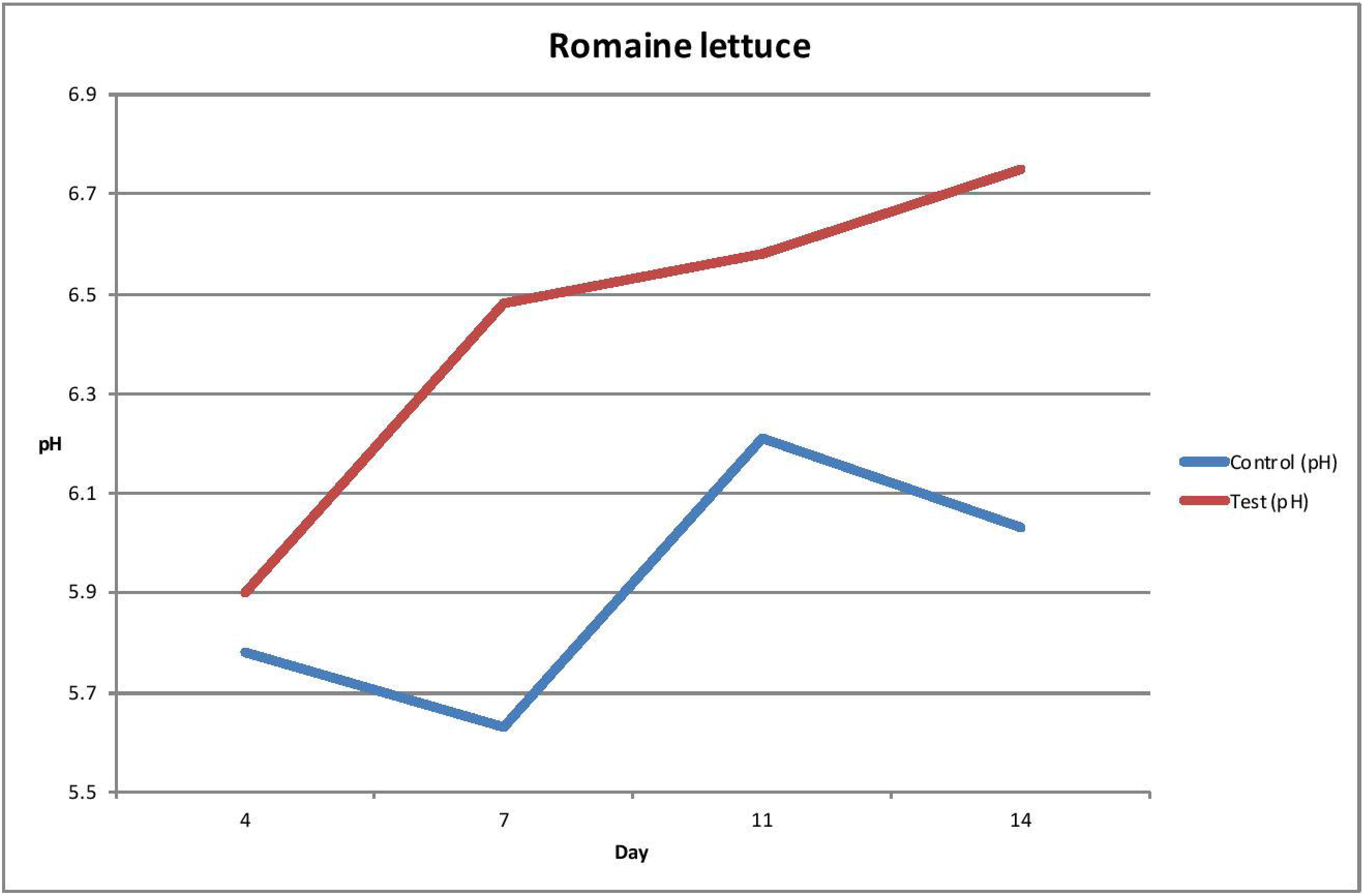
The results were not what we expected. The plant exposed to excess CO_2_ became more alkaline instead of more acidic, while the control plant remained mostly under pH=6.3. This suggest *Romaine lettuce* also has a homeostatic system to prevent over-absorption of CO_2_ in a CO_2_-rich environment.

### Results: *Nevada lettuce (Lactuca sativa)*

**Table 3:** pH readings vs exposure time (Days) in Nevada lettuce

| Day | Control<br>(pH) | Test (pH) |
| --- | --- | --- |
| 4 | 5.52 | 6.02 |
| 7 | 6.19 | 6.38 |
| 11 | 5.98 | 6.56 |
| 14 | 6.24 | 6.51 |

**Figure 3:**
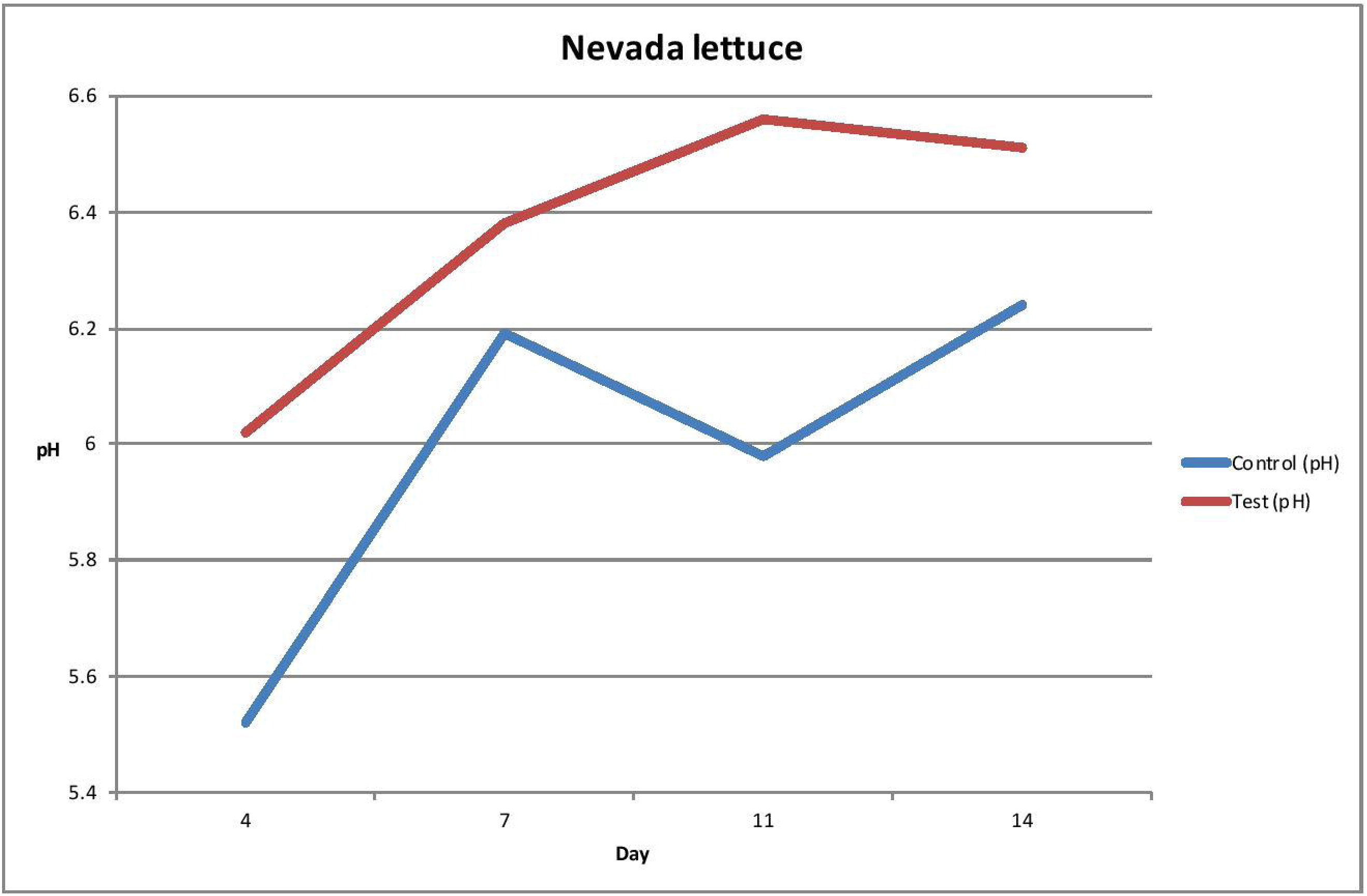
The results were not what we have expected. The plant exposed to excess CO_2_ became more alkaline instead of more acidic, while the control plant remained well under pH=6.3. This suggest *Nevada lettuce* also has a homeostatic system to prevent over-absorption of CO_2_ in a CO_2_-rich environment.

## Discussion and Conclusion

At the end of our experiment, we found pH levels moving in the opposite direction of what we had hypothesized. That is, the pH level of plants exposed to an increased level of CO_2_ increased, compared to the control plants. The more CO_2_ the plant was exposed to, the higher the pH of its internal environment became. We reject our initial hypothesis that a plant exposed to an increased CO_2_ will decrease the pH of its cytosol. Based on these results, we believe plant systems have unique homeostatic mechanism that drives out excess CO_2_ which results in the loss of normal levels of CO_2_ and causes the plant to run in a CO_2_ deficit. This would explain the increase in pH. This homeostatic system needs further investigation.

After performing a 2-Way ANOVA, we can conclude that statistically, there is no difference in the pH between the two different plants (p=1); but there is a difference in the pH between plants exposed to higher CO_2_ and normal CO_2_ (p=0.012); there is no interaction between type of plant and CO_2_ level (p=0.65).

### Further Hypotheses

We further hypothesize that the exposure of plants to increased CO_2_ accelerates the rate of plant glucose breakdown.

## Acknowledgment

We thank the Friends University VPAA office for the provision of starter funds to support research involving undergraduates.

