## Supplemental figure for "Investigating Increased CO_2_ concentration on the pH of various plant species"

Supplemental Information

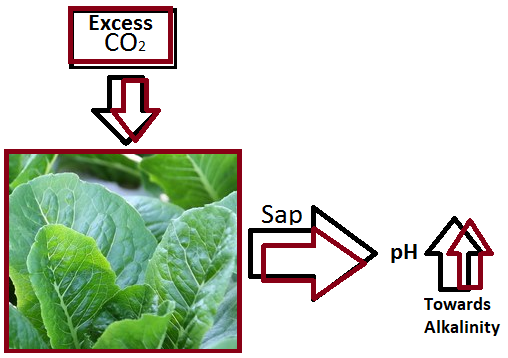

**Suppl. Figure 1:** Graphical Abstract

**Suppl Table 1***:* pH Data

|  | **TYPE OF PLANT** | | |
| --- | --- | --- | --- |
| **pH** | **Normal CO_2_** | **Romaine Lettuce** | **Nevada Lettuce** |
|  |  | 5.78 | 5.52 |
|  |  | 5.63 | 6.19 |
|  |  | 6.21 | 5.98 |
|  |  | 6.03 | 6.24 |
|  | **High CO_2_** | 5.9 | 6.02 |
|  |  | 6.48 | 6.38 |
|  |  | 6.58 | 6.56 |
|  |  | 6.75 | 6.51 |

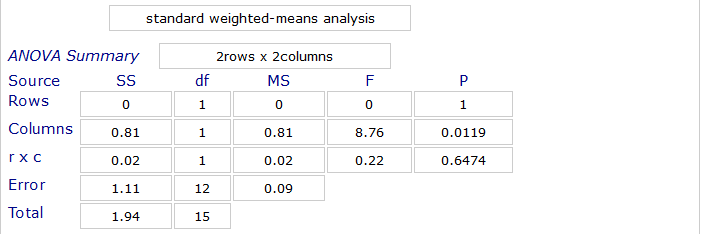

**Suppl. Figure 2**: 2-way Anova test results
